# The evolutionary advantage of condition-dependent recombination in a Red Queen model with diploid antagonists

**DOI:** 10.1101/478966

**Authors:** Sviatoslav R. Rybnikov, Zeev M. Frenkel, Tzion Fahima, Abraham B. Korol

## Abstract

Antagonistic interaction, like those between a host and its parasite, are known to cause oscillations in genetic structure of both species, usually referred to as Red Queen dynamics (RQD). The RQD is believed to be a plausible explanation for the evolution of sex/recombination, although numerous theoretical models showed that this may happen only under rather restricted parameter values (selection intensity, epistasis, etc.). Here, we consider two diploid antagonists, each with either two or three selected loci; the interaction is based on matching phenotypes model. We use the RQD, whenever it emerges in this system, as a substrate to examine the evolution of one recombination feature, condition dependence in diploids, which still remains an underexplored question. We consider several forms of condition-dependent recombination, with recombination rates in the host being sensitive either to the parasite’s mean fitness, or to the host’s infection status, or to the host’s genotype fitness. We show that all form of condition-dependent recombination can be favored over the corresponding optimal constant recombination rate, even including situations in which the optimal constant recombination rate is zero.

## BACKGROUND

Antagonistic interactions, like those between a host and its parasite, a prey and its predator, or a plant and its herbivore, are omnipresent in nature. Such interactions are long known to cause more or less regular oscillatory dynamics. In early 1920s Lotka [1] and Volterra [2] theoretically predicted oscillations in the *ecological* dynamics of the antagonists, i.e. in their population size or population density. The anticipated pattern was then observed both in field and lab; remarkably, pairs of host/parasite [3–5], prey/predator [6–8], and plant/herbivore [9] behaved in a qualitatively similar way, suggesting inherent commonality of underlying mechanisms. Later on, Haldane [10] hypothesized, preceding from general speculations on the negative frequency-dependent selection, that the antagonistic interactions should cause oscillations also in *genetic* dynamics of the antagonists, i.e. in allele frequencies of their interaction-related genes. This phenomenon is now known as ‘Red Queen dynamics’ (RQD), following the colorful terminology introduced by Van Vallen [11] and adopted by Bell [12]. Interestingly, RQD has become commonly discussed much prior to empirical tests for its existence [13,14], and even for the existence of negative frequency-dependent selection [15–18].

In late 1970s – early 1980s, several evolutionary biologists suggested, more or less explicitly, that the RQD may create preconditions for the evolution of sex/recombination in the host [19–24]. Following Bell [12], this assumption is now referred to as the ‘Red Queen hypothesis’ (RQH). It was successfully confirmed by the first formal model [25], but further simulations showed that the affirmative answer is much less general [26,27]. Reviewing numerous so far developed RQH-models allows inferring that antagonistic interactions favor sex/recombination under rather restrictive conditions. The ‘success’ depends on a number of factors, including type of genetic interactions between the antagonists [28,29], severity of the ‘antagonicity’ [30], strength of selection, at least in the parasite [31], epidemiological context [32], etc. Expectedly, the success is facilitated by imposing additional ‘pressure’ on the parasite, like similarity selection [33] or external abiotic selection [34]. Importantly, it is also clear now that in diploids, where sex implies both segregation and recombination, conditions favoring segregation *per se* (i.e. ‘sex without recombination’ versus ‘no sex’) and those favoring recombination *per se* (i.e. ‘sex with recombination’ versus ‘sex with no recombination’) are not the same [35,36]. In general, empirical studies do support the RQH, although the evidence remains very limited for sex [37–42] and even more so for recombination [43,44].

Here, we study the evolution of recombination under antagonistic interactions. However, we do not aim to further clarify in which situations antagonistic interactions result in the emergence of RQD. Instead, we test whether RQD, whenever it emerges, can favor one of the most intriguing features of recombination, its condition dependence (CD). By CD we understand the sensitivity of recombination rates (RRs) to external and/or internal conditions. This sensitivity can be manifested in at least two different ways. First, RRs can vary within *the same genotype* across its environment. Second, RRs can vary across *different genotypes* according to their fitness, which depends, in turn, on stress-tolerance or genetic background. Here, we refer to these two forms of CD recombination is environment-dependent (ED) and fitness-dependent (FD) recombination. To date, ED recombination has been reported for many species exposed to various environmental stressors (for recent reviews, see [45–47]). Yet, recombinogenic effect of antagonistic interactions remains obscure [48,49]. Empirical evidence for FD recombination is much more limited. The phenomenon was observed only in several experimental studies, where examined genotypes differed in their tolerance to extreme temperatures [50–53], desiccation [54], or toxic nutrients [55,56], or in the spectrum of deleterious mutations [57,58]. FD recombination under antagonistic interaction has never been reported, to the best of our knowledge.

The empirical evidence for CD recombination comes mostly from diploids. At the same time, theoretical diploid-selection models confirmed the evolutionary advantage of ED recombination [59,60], but obtained less clear results for FD recombination [60–62]. These theoretical studies considered cyclical selection and mutation–selection balance. Here, we test whether CD recombination can be evolutionarily advantageous in a diploid host involved in RQD with its diploid parasite. Earlier, it was found that RQD may create preconditions for the evolution of CD sex [63]. Here, we focus on recombination *per se*, assuming obligate sex in both antagonists. In both antagonists, we assume either two or three selected loci. ED recombination is examined in both models, while FD recombination – only in the model with three selected loci (the reason is that recombination matters only in genotypes heterozygous for at least two loci, and in models with two selected loci there exists only one such genotypes, which means no variation in fitness). We test whether or not CD recombination is evolutionary advantageous by comparing it with the optimal constant RR. At that, such comparisons are based on allele dynamics at a selectively neutral locus affecting RRs only (recombination modifier).

## MODEL AND METHODS

### Life cycle

Both species reproduce sexually, by outcrossing, with total panmixia. The populations are infinite. The generations do not overlap. In both species, life cycle includes fertilisation, maturation (selection), meiosis (recombination), and random mating. Let *x*_*ij*_ be zygote/adult of genotype *ij* (*i* and *j* stand for the parental haplotypes), and *g*_*k*_ be gamete of haplotype *k*. The frequency *p’* of genotype *x*_*ij*_ after selection (adults) is proportional to frequency *p* of this genotype before selection (zygotes), and to its absolute fitness *W*:

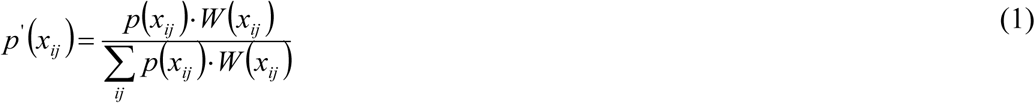

The adults produce gametes; frequency of gamete *g*_*k*_ in the gamete pool is:

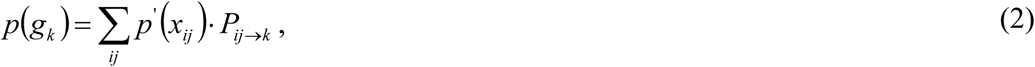

where *P*_*ij*→*k*_ is the probability of obtaining gamete g_*k*_ from genotype *x*_*ij*_ [64]. The gametes form zygotes of the next generation; given the random mating, frequency of zygote *x*_*ij*_ is proportional to frequencies of both parental gametes:

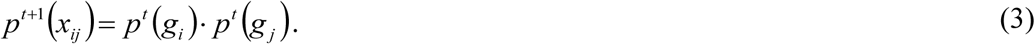

### Selected system and selection regime

In both species, the selected system consists of either two or three linked interaction-mediating loci. At each selected locus *l*, there exist two possible alleles: 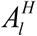 and 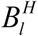 in the host, and 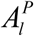 and 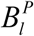 in the parasite. The host’s homozygotes 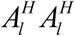 and 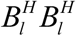 display phenotypes 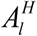 and 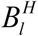, respectively, while the parasite’s homozygotes 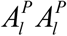 and 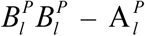 and 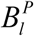, respectively. In both species, one of the alleles at each selected locus is fully dominant in relation to the other, so that heterozygotes display phenotype equal to that of one of the homozygotes. The encounter between a given host, *H*_*i*_, and a given parasite, *P*_*j*_, results in one of the two possible outcomes, either resistance or infection, depending on their phenotypes. If there exists at least one selected locus *l* at which the host’s and the parasite’s phenotypes differ (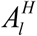 and 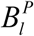, or 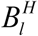 and 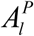), then the host ‘recognizes’ the parasite and demonstrates resistance; otherwise, infection emerges. We refer to such type of interaction as ‘matching-phenotype model’, which can be viewed as an extension to diploids of the canonical haploid matching-genotype models [21,25,65]. Both host fitness, *ω*^*H*^, and parasite fitness, *ω*^*P*^, are determined by the infection/resistance status. In the host, resistance means unit fitness, while infection reduces it by a certain value, *s*^*H*^. Vice versa, in the parasite, infection means unit fitness, while resistance reduces it by a certain value, *s*^*P*^:

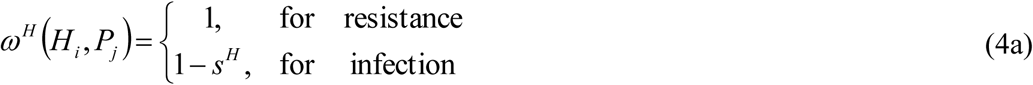

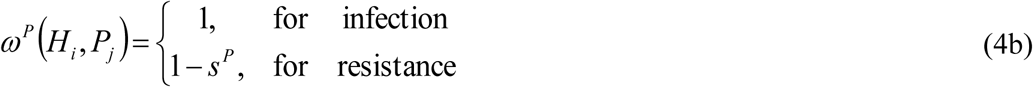

Therefore, the two parameters, *s*^*P*^ and *s*^*E*^, characterize selection intensity: higher values mean weaker selection and vice versa.

Both species are subject to frequency-dependent selection. The frequency of encounters between a given host, *H*_*i*_, and a given parasite, *P*_*j*_, is proportional to their frequencies in the corresponding populations, *p*(*H*_*i*_) and *p*(*P*_*j*_). Then, absolute fitness of the considered host class, *W*(*H*_*i*_), can be calculated as the expected values of *ω*^*H*^(*H*_*i*_, *P*_*j*_), and absolute fitness of the considered parasite class, *W*(*P*_*j*_), – as the expected values of *ω*^*P*^(*H*_*i*_, *P*_*j*_):

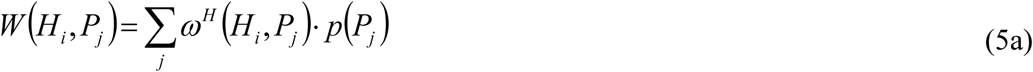

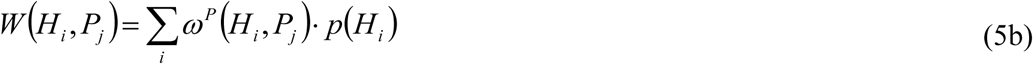

### Recombination strategies

In the considered model, RRs within the selected system of the parasite were fixed and, therefore, treated as a parameter. In the host, RRs within the selected system were determined by genotype at an additional selectively neutral locus (recombination modifier). In both species, in three-locus selected system, RRs between both pairs of the adjacent selected loci were for simplicity assumed to be equal. In the host, the modifier locus was assumed to be unlinked to the selected system.

In the host, modifier alleles conferred either constant recombination (with RRs equal for all hosts and stable in time), or CD recombination (with RRs varying among the hosts). We considered three forms of CD recombination: (*i*) parasite-dependent (PD) – with RRs varying among the hosts according to the parasite mean fitness; (*ii*) infection-dependent (ID) – with RRs varying among the hosts according to their infection status; and (*iii*) fitness-dependent (FD) – with RRs varying among the hosts according to their own fitness. Under all forms of CD recombination, RRs negatively depended on conditions: ‘bad’ conditions (measured as either infection, or high fitness of the parasite, or low fitness of the host) caused higher RRs and vice versa. PD and ID recombination were examined in the models with two and three selected loci, while FD recombination – only in the model with three selected loci. The reason is that FD recombination is imaginable only in the presence of variation in fitness among those genotypes in which recombination has fitness consequences in the next generation, which requires at least three selected loci [60,62].

CD recombination was compared with the corresponding (for the given parameter set) optimal constant RR. In situations with *zero* optimal constant RR, the competing CD recombination implied an increase in RR in ‘bad’ conditions (recombination-increasing strategy). In situations with *non-zero* (intermediate) optimal constant RR, three types of CD strategies were considered, differing in their effect on RR: (*i*) recombination-increasing – only with an increase in RR in ‘bad’ conditions; (*ii*) recombination-decreasing – only with a decrease in RR in ‘good’ conditions; and (*iii*) ‘fringe’ – with an increase in RR in ‘bad’ conditions and a decrease in RR in ‘good’ conditions.

### Comparison of recombination strategies

The examined recombination strategies were compared within the modifier approach [66,67], based on dynamics of the corresponding recombination-modifier alleles. One modifier allele was regarded as capable to invade the population, which currently occupied by the alternative modifier allele, if the former increased its frequency upon injection with initial frequency 5%. We classified the outcomes of competition between CD recombination and the optional constant RR as follows: (*i*) disadvantage of CD recombination – its allele could not invade the population occupied by allele for the optimal constant RR; (*ii*) partial advantage – both allele for CD recombination and allele for the optimal constant RR could invade the population occupied by the competing allele; (*iii*) full advantage of CD recombination – its allele could invade the population occupied by allele for the optimal constant RR, while the latter could not invade the population occupied by the former. In all pairwise comparisons, the modifier alleles were traced during 1,000 generations, with a ‘burn-in’ period of 4,000 generations needed for the system to enter the oscillation regime. Naturally, a constant RR was regarded as optimal if it demonstrated full advantage over both lower and higher RRs. The interaction between all recombination-modifier alleles was assumed to be purely co-dominant.

## RESULTS AND DISCUSSION

### Conditions for non-damping oscillations and selection for/against non-zero RRs

We considered a variety of selection regimes, differing in selection intensity in the host and in the parasite, *s*^*H*^ and *s*^*P*^, respectively. These parameters were scanned in the range from 0.05 (mildly deleterious infection) to 1 (lethal infection). The third scanned parameter was RR in the parasite, *r*^*P*^. Naturally, studying the evolution of recombination requires polymorphism maintenance in the selected system. Conditions compatible with polymorphism maintenance in models with diploid antagonists are formulated only partially, especially for multilocus genetic systems [68–71]. For the considered type of interaction, we observed protected polymorphism only if both antagonists had the same dominance/recessiveness relations at their selected loci, i.e., if the host’s alleles 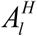 and the parasite’s alleles 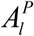 were either simultaneously dominant or simultaneously recessive; a situation was regarded as polymorphic if along the trajectory each selected allele had frequency >10^−4^. However, the context of our study required not only polymorphism maintenance in the selected system, but also stable RQD. Under most tested parameter sets, the oscillations appeared to be damping, although the decay was often rather slow; oscillations were regarded as damping if the amplitudes of population mean fitness in both species consequentially decreased during 10,000 generations.

We examined situations with both zero and non-zero (intermediate) optimal constant RR within the selected system of the host. Zero optimal constant RR was favored in a wide area of the parameter space; however, this often happened under no oscillations, or with oscillations damping in time. The demand of non-dumping RQD, with *zero* optimal constant RR in the host, was met under weak selection and very low RR in the parasite (*s*^*P*^ ≤ 0.34 and *r*^*P*^ ≤ 0.001 in the model with two selected loci, *s*^*P*^ ≤ 0.40 and *r*^*P*^ ≤ 0.003 in the model with three selected loci). In the model with two selected loci, this demand also required weak selection in the host (*s*^*H*^ ≤ 0.24), while in the model with three selected loci this parameter appeared to be not crucial (*s*^*H*^ ≤ 0.95). Non-dumping RQD with *non-zero* optimal constant RR in the host was found only in the model with two selected loci. This happened under strong selection and low RR in the parasite (*s^P^* ≥ 0.76, *r*^*P*^ ≤ 0.004), and also under not too strong selection in the host (*s*^*H*^ ≤ 0.84). At that, restrictions on the selection intensity in the host became more stringent as selection in the parasite weakened (0.10 ≤ *s*^*H*^ ≤ 0.84 for *s*^*P*^ = 1.00, compared to 0.14 ≤ *s*^*H*^ ≤ 0.16 for *s*^*P*^ = 0.76). The optimal constant RR in the host varied within almost the whole range of possible values (0.013 ≤ *r*^*P*^ ≤ 0.469). The optimal constant RR in the host appeared to be in strong positive correlation with selection intensity in the host, and with RR in the parasite. In the model with three selected loci, we did found several cases with very low but still non-zero optimal constant RR (*r*^*H*^ ≤ 0.004), and with non-damping oscillations in mean fitness of the populations; yet, the amplitude of the oscillations was too small (<0.1%) to consider such cases as a stable biologically interesting RQD. The low optimal RRs in the host are in line with the results obtained in some other RQD models [65,72–74]. Potentially, selection for non-zero RR in the host could be facilitated by assuming other forces, like similarity selection [33], which requires further exploration.

### The evolutionary advantage of CD recombination in the host

Our simulations confirmed that all three examined forms of CD recombination (PD, ID and FD recombination) can be favored over the corresponding optimal constant RR, including situations in which the optimal constant RR is zero (Fig. 1, 2). With zero optimal constant RR, cases favoring CD recombination were usually characterized by stronger selection and larger amplitude of oscillations, compared to cases rejecting CD recombination. At that, PD recombination tended to require higher selection intensity in the parasite, while ID recombination – in the host. In contrast, with non-zero optimal constant RR, cases favoring CD recombination usually had weaker selection.

**Fig. 1.**
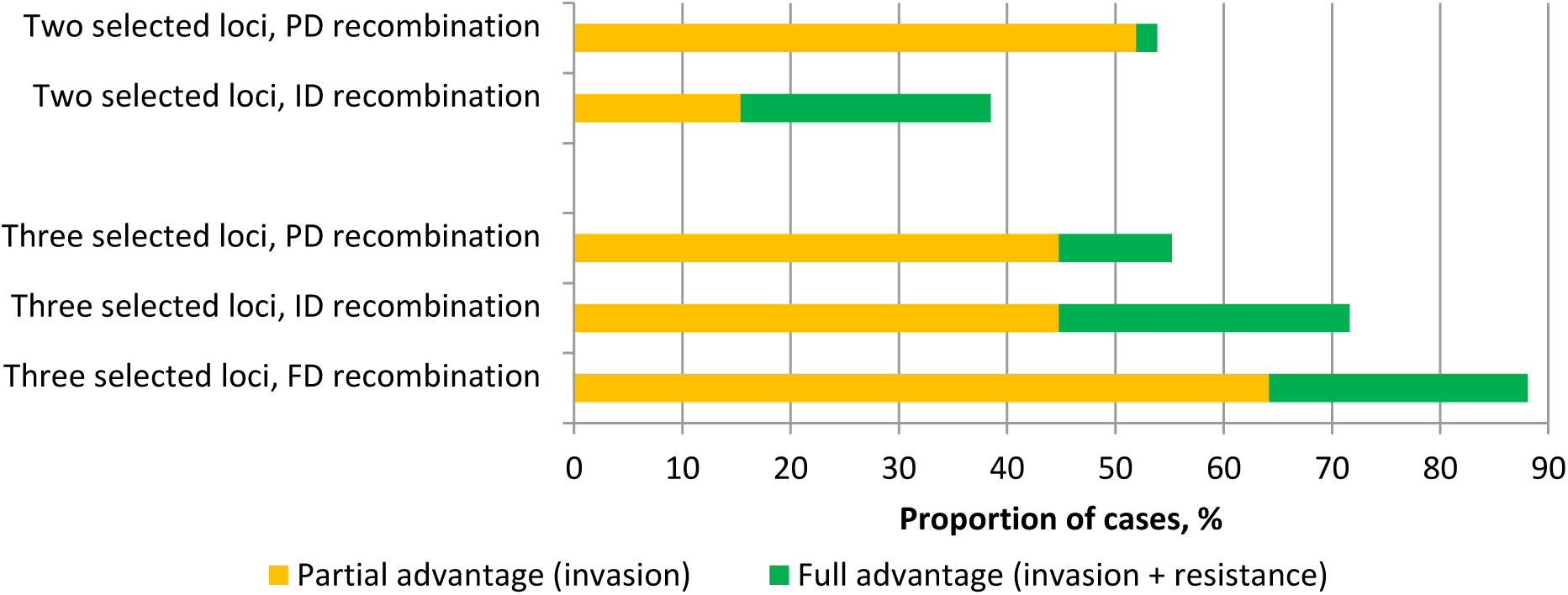
Proportion of cases in which different forms of CD recombination are favored over the corresponding zero optimal constant RR (the models with two- and three selected loci)

**Fig. 2.**
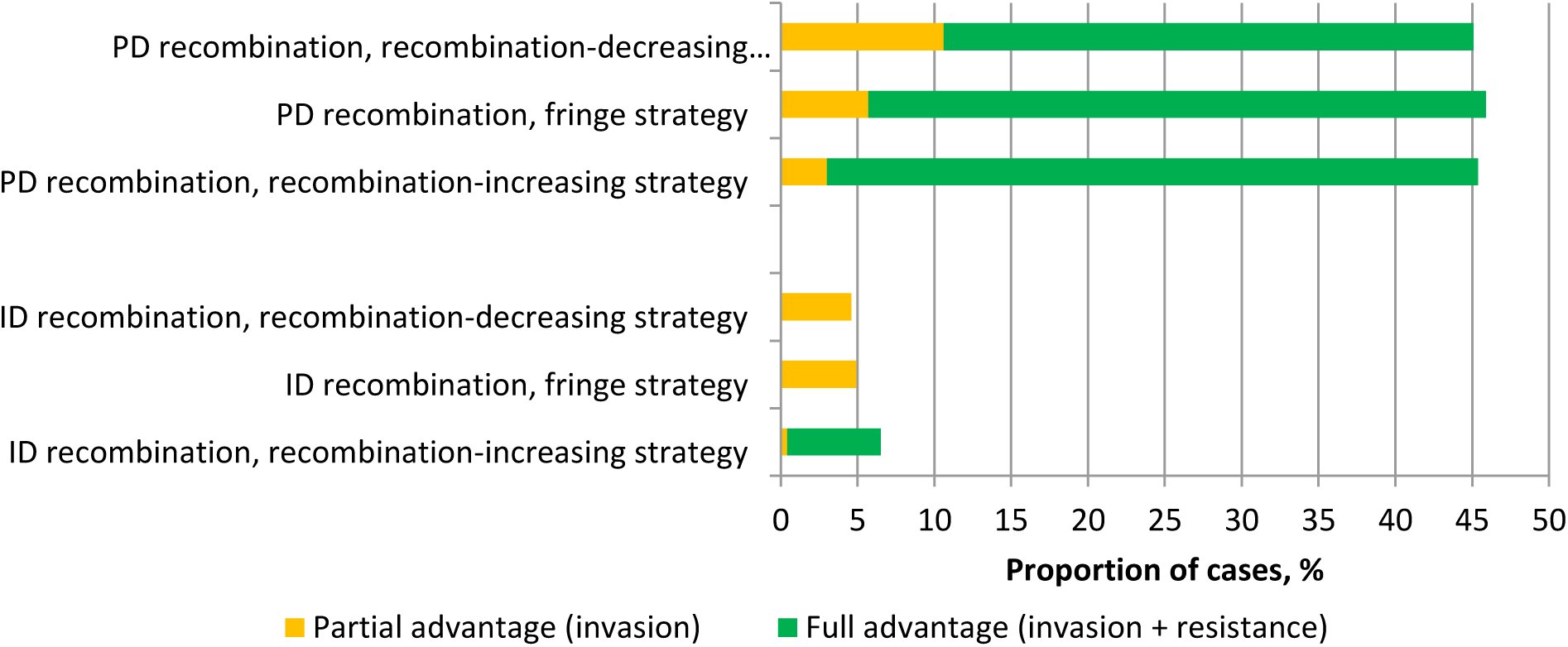
Proportion of cases in which different forms of CD recombination are favored over the corresponding non-zero (intermediate) optimal constant RR (the model with two selected loci)

A common explanation for the evolutionary advantage of CD recombination is the so-called ‘abandon-ship’ mechanism, appealing to benefits obtained by a recombination-modifier allele from adjusting its own linkage to the selected system [61]. Another explanation is benefits of RR plasticity within the selected system, which arise due to a ‘protection’ of better allele combinations. Here, we intentionally assumed recombination-modifier locus to be unlinked to the selected system, in order to exclude any selfish benefits. Thus, wherever CD recombination was favored in our models, this must be ascribed to benefits of such strategy for the selected system. It is noteworthy that mean RR in the population practicing CD recombination may deviate from the optimal constant RR (in case of ‘one-sided’ strategies, such as recombination-decreasing and recombination-increasing ones, this happens inevitably). Thus, whether CD recombination will or will not be favored over the corresponding optimal constant RR, depends on which of the two effects, the benefits of RR plasticity or the costs of deviation from the optimal RR, outbalances in a given situation. At that, both the benefits and the costs are function of amplitude of RR plasticity. The interplay between these two function is not trivial, and the net function can be even non-monotonous, so that an intermediate amplitude of RR plasticity sometimes appears optimal, as in our previous study [62].

The examined forms of CD recombination appeared to be favored over the corresponding optimal constant RR in different proportion of cases. In the model with three selected loci (in which all three forms of CD recombination are imaginable), FD recombination was the most advantageous. This seems reasonable since two other forms (PD and ID recombination) do not take into account the difference between the host genotypes (although this difference does exist in systems with three selected loci) and, therefore, reflect only ecological plasticity of RR. In contrast, FD recombination includes sensitivity to variation in fitness (explicitly, by definition) and environment (implicitly, given the fact that fitness of genotypic classes of the host depends on frequencies of all genotypic classes of the parasite). The difference between PD and ID recombination may arise from peculiarities of the emerged RQD patterns. For example, in the model with two selected loci, PD recombination was often favored over the corresponding intermediate optimal RR, whereas ID recombination was not. It turned out that in such situations the proportion of infected heterozygotes in the host population oscillates with about twice longer period than mean fitness in the parasite population, thereby “buffering the input information” on the parasite for the host (Fig. 3). As a result, ID recombination, which implies sensitivity to the host infection status, “missed” a significant part of information on the environment.

**Fig. 3.**
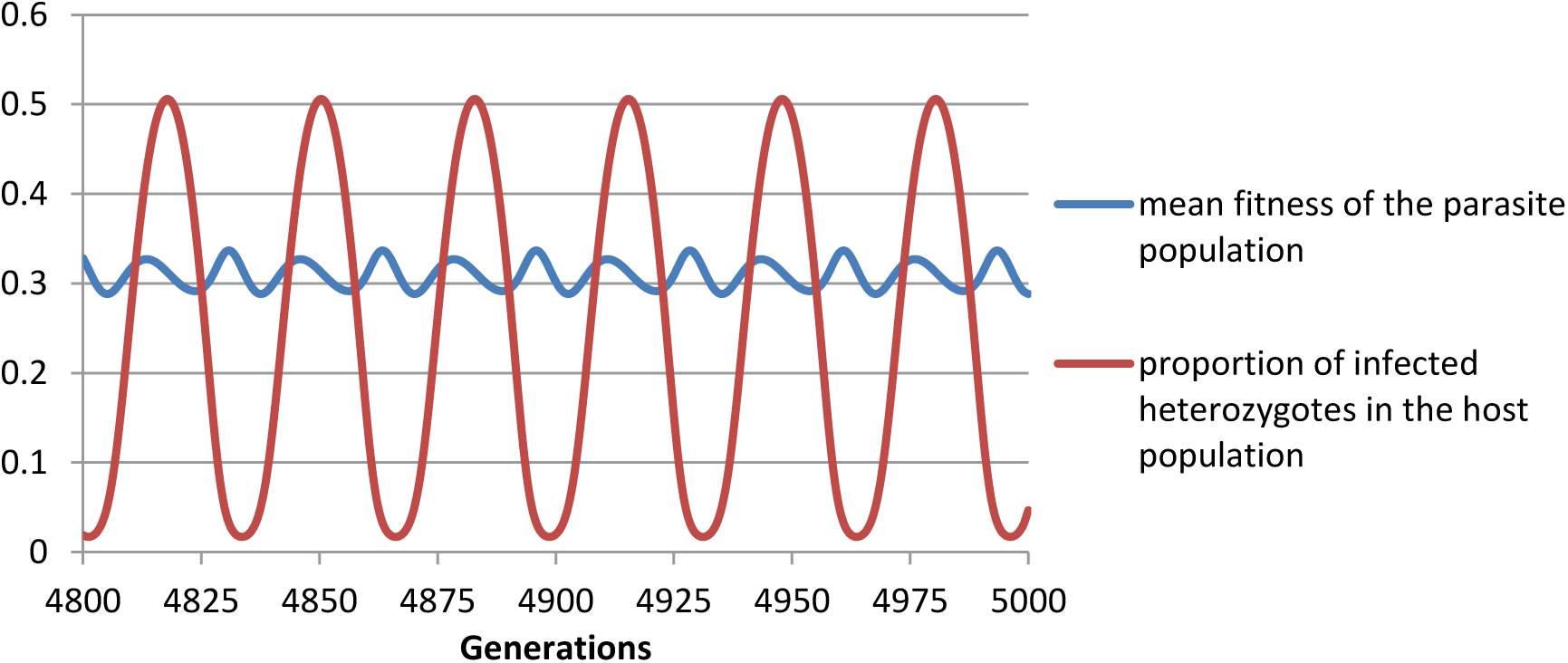
An example of RQD where mean fitness of the parasite population and proportion of infected heterozygotes in the host population oscillate with different periods (the model with two selected loci, *s*^*P*^ = 0.92, *s*^*H*^ = 0.22, *r*^*P*^ = 0.001, *r*^*H*^ = 0.081)

In addition to the above presented matching-phenotype model, we also examined a quantitative-trait model [68]. We considered situations where each of the antagonists had three selected loci which together encoded an interaction-mediated trait. In both species, selection intensity was assumed to be a power function of the absolute difference between the host’s and the parasite’s trait values. In this model, situations favoring non-zero RRs in the host were observed very rarely. Typically, they required strong selection in the parasite, weak to intermediate selection in the host, and very low RRs in the parasites. Weak selection in both antagonists usually resulted in rejection of non-zero RRs in the host. We found situations with in which FD recombination was favored over the corresponding optimal constant RRs in the host, both zero and non-zero. In cases with zero optimal constant RR, the evolutionary advantage of FD recombination tended to decrease with higher amplitude of RR plasticity, confirming the negative effect of deviation from the optimal RR on the evolution of CD recombination.

